# The combination of *EWSR1-FLI1* and loss of one *EWSR1* allele leads to the induction of trisomy 8

**DOI:** 10.64898/2026.05.21.726567

**Authors:** Harsha Hapugaswatta, Alejandro Parrales, Hyewon Park, Haeyoung Kim, Tomoo Iwakuma, Mizuki Azuma

## Abstract

Ewing sarcoma is a pediatric cancer that develops in skeletal elements. The majority of Ewing sarcoma patients carry the aberrant *EWSR1-FLI1* fusion gene. Despite trisomy 8 being an additional common aberration associated with a poor prognosis for patients, its induction mechanism remains unknown. When the *EWSR1-FLI1* gene is formed, the cell loses one wildtype EWSR1 allele. To elucidate the induction mechanism of trisomy 8, we generated a cell line that allows for the conditional induction of EWSR1-FLI1 expression and EWSR1 knockdown (derived from a single *EWSR1* allele. Specifically, the conditional cell line was generated by integrating the Tet-on *EWSR1-FLI1* construct into the *AAVS* locus and adding a *miniAID* tag at the 5’ end of the EWSR1 locus using auxin-degron system. A combination of the EWSR1-FLI1 expression and degradation of one allele-derived EWSR1 induced a high incidence of trisomy 8 within eight days, enhancing colony formation. Mechanistically, trisomy 8 is induced by the haploinsufficiency of EWSR1, and the remaining EWSR1 proteins are likely inhibited by interaction with EWSR1-FLI1. Our data showed that the knockout of EWSR1 alone was sufficient to increase the incidence of trisomy 8. Expression of wild-type EWSR1 in EWSR1 knockout cells rescued the high incidence of trisomy 8. In contrast, the EWSR1:R565A mutant, which lacks the ability to interact with Aurora B kinase, failed to rescue this phenotype. We propose that the combination of EWSR1-FLI1 expression and loss of EWSR1 contributes to the induction of trisomy 8 through the compromised EWSR1-Aurora B pathway.

**Graphical Abstract:** 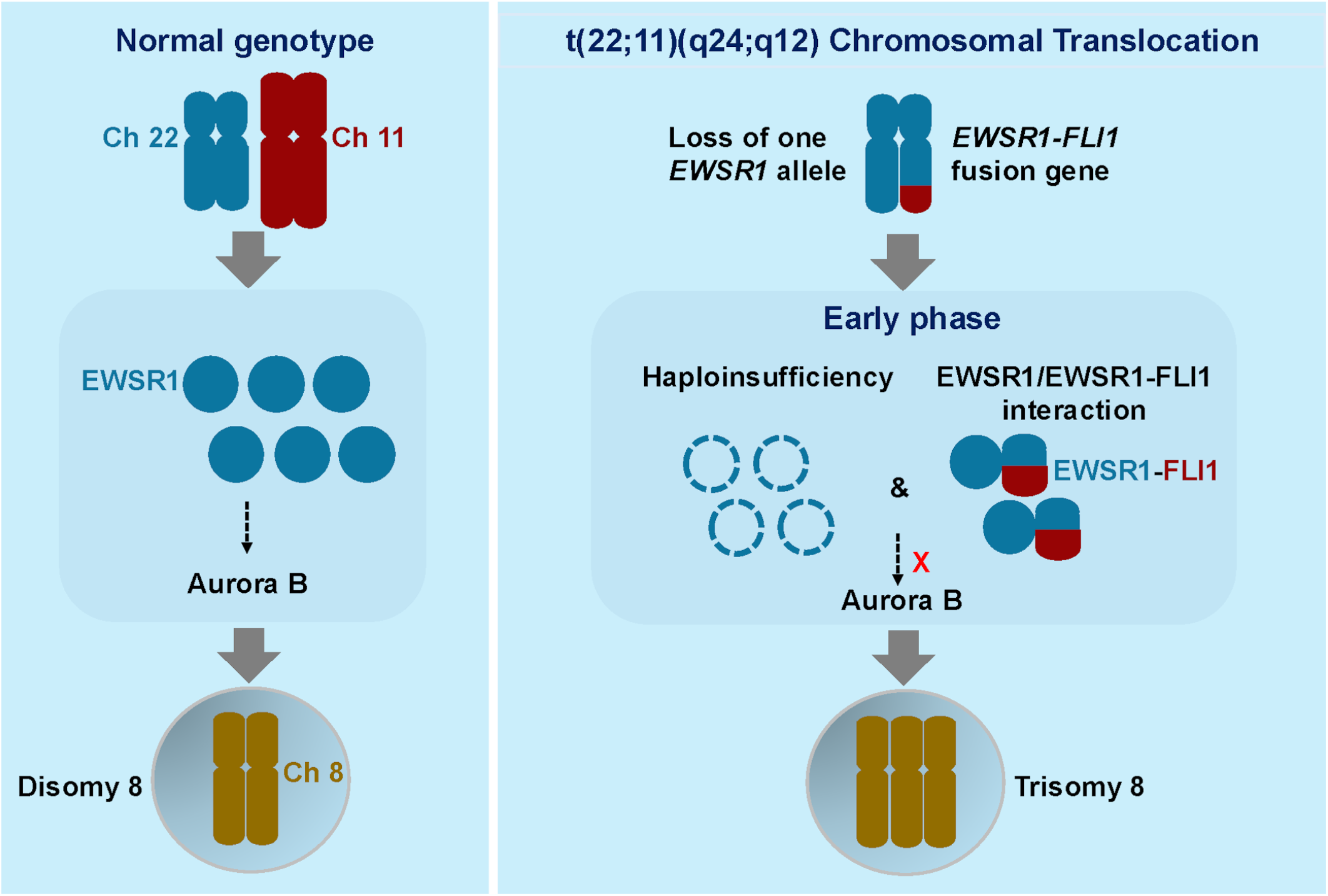

## Introduction

Ewing sarcoma is the second most common type of bone cancer that develops in children and adolescents. Although the survival rate for patients with primary tumors has improved to approximately 70% in recent decades, the remaining patients experience relapse and/or metastasis with a low survival rate of less than 10% (*1, 2*). To target cells with relapse and/or metastasis, it is necessary to understand the underlying mechanism. All Ewing sarcoma cells carry a fusion gene comprising the N-terminus of *EWSR1* and the C-terminus of one of the *ETS* genes (*FLI1, ERG, ETV1, ETV4, and FEV*) (*3-7*). Relapse and/or metastasis often involve acquisition of additional secondary aberrations, and its examples are aneuploidy (changes in DNA copy numbers), DNA mutations, epigenetic changes, and RNA mis-splicing (*8-14*). Trisomy 8, which is a condition when a cell acquires three copies of chromosome 8, is present in approximately 50% of Ewing sarcoma patients (*15-17*). Trisomy 12 is another aneuploidy detected in 20% of patients, while a gain of 1q is observed in approximately 25% of patients (*17, 18*). Among these aneuploidies, trisomy 8 is particularly associated with a poor prognosis. Supporting this observation, the *RAD21* gene, encoded on chromosome 8, is overexpressed when the cell gains the third chromosome 8. Its overexpression mitigates replication stress and provides a survival advantage (*16*). Chromosome 8 also encodes a number of genes that drive tumorigenesis, including *c-MYC* and *FGF Receptor (FGFR*), potentially contributing to the progression of the disease (*19, 20*). While these observations highlight the importance of trisomy 8 in disease progression, the mechanism by which it is induced remains unclear. Therefore, we aimed to gain insight into the induction mechanism of trisomy 8 by studying Ewing sarcoma-associated proteins.

In a previous study, we demonstrated that EWSR1 plays a crucial role in maintaining faithful mitosis and preventing the induction of aneuploidy by regulating the key mitotic kinase, Aurora B (*21-24*). The Aurora B kinase is a crucial player in ensuring the accurate segregation of chromosomes, resulting in the equal distribution of chromosomes into two daughter cells (*25, 26*). Compromised chromosomal segregation can induce aneuploidy (*27, 28*). Our study revealed that EWSR1 regulates mitosis by physically interacting with Aurora B (*22*). The arginine (Arg) located at the 565^th^ amino acid of EWSR1 is essential for its interaction with Aurora B kinase, thereby preventing the induction of aneuploidy. It is important to note that the EWSR1-FLI1 does not contain this amino acid, as it only includes the N-terminus of the protein. Furthermore, the importance of maintaining chromosomal integrity in suppressing tumorigenesis was highlighted in our previous study using the zebrafish *ewsr1a* (a homologue of human *EWSR1*) mutant (*23*). The study revealed that homozygous and heterozygous *ewsr1a* mutants spontaneously develop a higher incidence of tumorigenesis compared to wildtype in the *tp53* mutant background. The study also demonstrated that the zebrafish *ewsr1a* homozygous mutant induces a high incidence of mitotic dysfunction and aneuploidy. Thus, the function of EWSR1 and Ewsr1a in maintaining chromosomal integrity is conserved between the two species.

Interestingly, mitotic phenotypes caused by the expression of EWSR1-FLI1 and EWSR1 knockdown are similar across multiple biological processes. An example of this is the induction of R-loops and the inhibition of BRCA1-dependent homologous recombination (*29*). Our previous study showed that both EWSR1-FLI1 expression and EWSR1 knockdown lead to an increase in the incidence of aneuploidy respectively (*23, 24, 30*). We demonstrated that the EWSR1-FLI1 phosphomimetic mutant with a Thr79 substitution to Asp (EWSR1-FLI1:T79D) causes a high incidence of aneuploidy, whereas a point mutation of Thr79 substituted for Ala in EWSR1-FLI1 (EWSR1-FLI1:T79A) fails to induce aneuploidy (*30*). Based on previous reports demonstrating the physical interaction between EWSR1-FLI1 and EWSR1, the phenotypic resemblance between expression of EWSR1-FLI1 and EWSR1 knockdown can be explained by the dominant inhibition of EWSR1 (*31-33*).

Based on our previous observations, we investigated the mechanism behind the induction of trisomy 8, the second most common DNA aberration observed in Ewing sarcoma. In this study, we demonstrate that trisomy 8 can be induced by the combination of EWSR1-FLI1 expression and one allele-derived EWSR1. Finally, we demonstrate that the EWSR1-Aurora B interaction is necessary to prevent the induction of trisomy 8. This is the first demonstration of factors contributing to the induction of trisomy 8.

## Results

### The combination of one allele-derived EWSR1 and EWSR1-FLI1 expression promotes colony formation

Approximately 50% of Ewing sarcoma patients carry trisomy 8, while 21% of patients carry trisomy 12 (*34*). Due to the frequent occurrence of trisomy 8, our aim was to identify the factors that induce these aneuploidies. Since the establishment of *EWSR1-FLI1* fusion gene is accompanied by the loss of one *EWSR1* allele, we aimed to systematically measure the induction of trisomy 8 upon the loss of one *EWSR1* allele, *EWSR1-FLI1*, or a combination of both. To achieve this, we edited the genome of DLD-1 cells using the CRISPR/Cas9 system, and established a conditional cell line capable of inducing these aberrations (**Fig 1A**) (*35*). The cells were engineered with three constructs: 1) *OSTIR1*, a plant-derived E3 ligase with a myc tag (10 amino acid) at the *RCC1* locus, 2) *miniAID*, a peptide with a plant-derived ubiquitination site for OSTIR1, fused with a FLAG tag at the 5’ end of one *EWSR1* gene, and 3) insertion of a Tet-on construct containing *EWSR1-FLI1-mCherry* at the *AAVS1* locus (*36-38*). We chose DLD-1 cell line for this study because it has a near-diploid karyotype without expression of EWSR1-FLI1, and it has been used to model changes in ploidy (*39, 40*). It is important to note that Ewing sarcoma cells are not ideal for studying the induction of trisomy 8, as they already carry this specific aneuploidy. Throughout the rest of the study, the cell line will be referred to as (*AID-EWSR1/wt;EWSR1-FLI1/wt*) cells. This cell line allowed us to systematically compare the activities of the one allele-derived EWSR1 (auxin (AUX) treatment), EWSR1-FLI1 expression (doxycycline (DOX) treatment), and their combination (AUX and DOX treatment) to the control (DMSO treated) cells.

**Fig 1.**
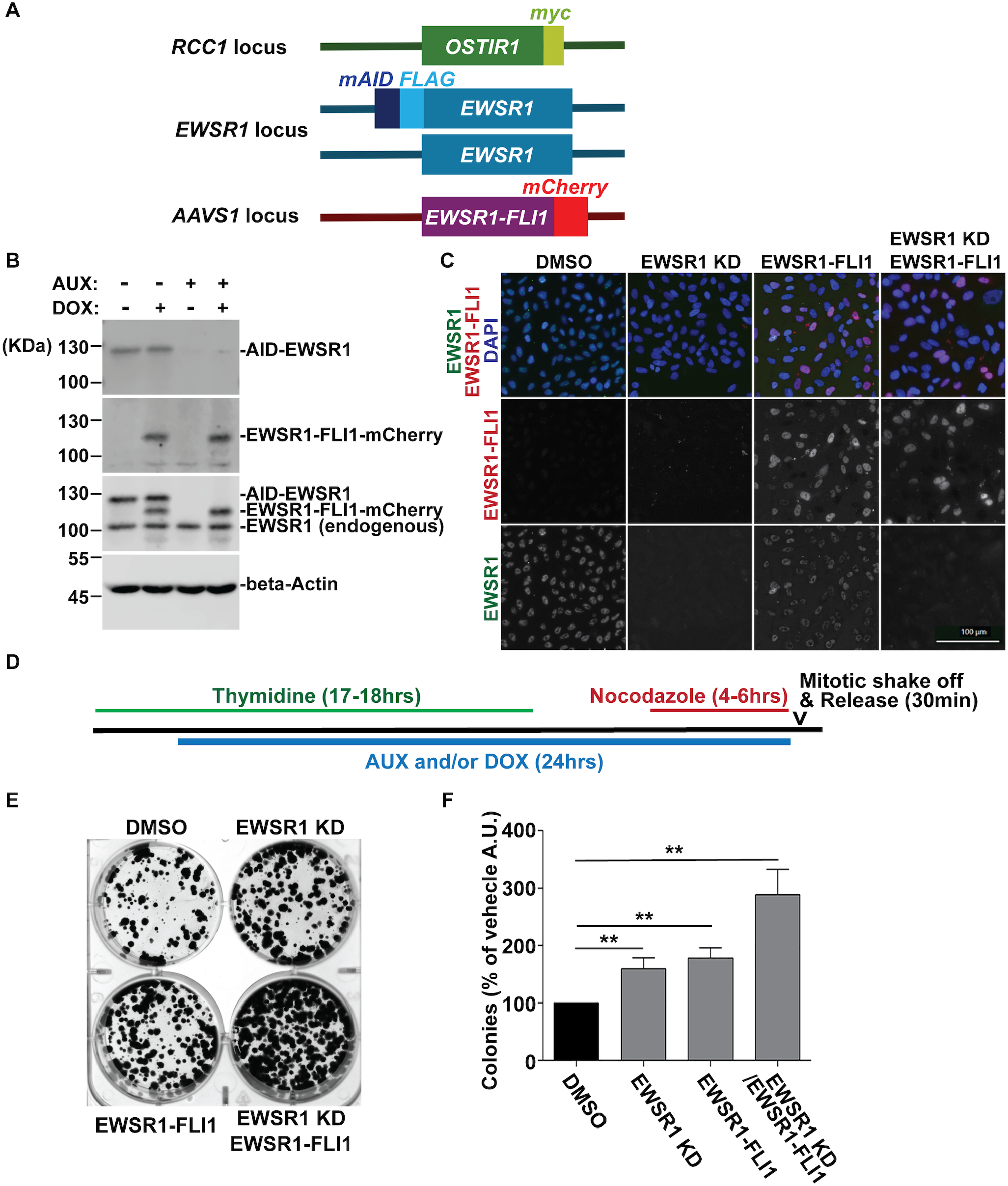
Combination of EWSR1 knockdown and EWSR1-FLI1 expression leads to high colony forming activity. **A**. Schematic diagram for generating the (*AID-EWSR1/wt;EWSR1-FLI1*) cell line. The plant E3 ligase, *OSTIR1*, was integrated into the *RCC1* locus. *mAID* tag was inserted into the *EWSR1* allele. The tet-on *EWSR1-FLI1* DNA construct was integrated into the *AAVS1* locus. **B**. Representative images of western blotting with anti-FLAG antibody (top panel), anti-mCherry antibody (second panel from the top), anti-EWSR1 (second panel from the bottom), and beta-Actin antibody (bottom panel). **C**. Representative images of immunocytochemistry. Top panel: merged images with anti-mCherry (Red), anti-FLAG (Green) and DAPI (Blue). Middle panel: anti-mCherry (Red). Bottom panel: anti-FLAG (Green) Scale bar: 100 µm. **D**. Schematic diagram for the mitotic synchronization, followed by colony formation assay. **E**. Representative photo images of the colonies assessed by colony formation assay. **F**. Numbers of the average numbers of colonies obtained from n=3 experiments. EWSR1 KD:EWSR1 Knockdown. ***: P<0.001, ****: P<0.0001.

To confirm the conditional degradation of mAID-EWSR1 protein derived from one *EWSR1* allele and the conditional expression of EWSR1-FLI1, the cell line was treated with AUX, DOX, AUX/DOX and, DMSO (control) for twenty-four hours. The western blotting confirmed the efficient degradation of mAID-EWSR1 in the AUX-treated cells and expression of EWSR1-FLI1 in the DOX-treated cells (n=3 independent experiments) (**Fig 1B**). In addition, the levels of mAID-EWSR1 in the AUX- and AUX/DOX-treated cells were lower than in the DMSO treated cells. Furthermore, the levels of EWSR1-FLI1 were significantly higher in the DOX and AUX/DOX treated cells compared to the control group (**Fig 1B**). The expression patterns of EWSR1 and EWSR1-FLI1 in the AUX- and/or DOX-treated cells were confirmed through immunocytochemistry using anti-FLAG for visualizing EWSR1 detection and anti-mCherry for EWSR1-FLI1 expression (**Fig 1C**). In the following sections, we will refer DMSO-treated cells as the control group, AUX-treated cells as EWSR1 knockdown (KD), DOX-treated cells as EWSR1-FLI1, and AUX- and DOX-treated cells as EWSR1 KD and EWSR1-FLI1 respectively.

To determine whether the one allele-derived EWSR1 and EWSR1-FLI1 expression enhances malignant properties, four samples were treated with DMSO, AUX, DOX, and AUX/DOX. These samples were then subjected to colony formation assays to measure the ability of a cell to grow three-dimensionally and form a colony. While AUX- and DOX-treated cells showed slight increase in colony-forming potential compared to the control (DMSO), the AUX/DOX double-treated EWSR1 KD/EWSR1-FLI1 cells displayed significantly higher colony forming potential (**Fig 1E and 1F**). This result suggests that the one allele-derived EWSR1 and the EWSR1-FLI1 expression cooperatively increased the colony-forming potential.

### The combination of EWSR1-FLI1 expression and one allele-derived EWSR1 promotes the induction of trisomy 8

Given that trisomy 8 is the most common aneuploidy observed in Ewing sarcoma, we next evaluated whether EWSR1-FLI1 and/or one allele-derived EWSR1 contribute to the induction of this aneuploidy. The cells were treated with DMSO, AUX, DOX, and AUX/DOX for two and eight days, followed by Fluorescence In Situ Hybridization (FISH) using a probe for chromosome 8 (**Fig 2A**). The number of signals per cell obtained from the probe was scored, and the percentages of cells with each number of signals were plotted (**Fig 2B and 2C**). The cells treated with AUX and/or DOX for two days did not display significant change in the rate of trisomy 8 (**Fig 2B**). Contrary, the treatment with AUX and DOX for eight days significantly increased the ploidy of chromosome 8 compared to the control cells. Specifically, trisomy 8 was more prevalent in the AUX/DOX treated cells (**Fig 2C**). These results suggest that the combination of one allele-derived EWSR1 loss and EWSR1-FLI1 expression leads to the highest incidence of trisomy 8, highlighting the importance of the additive activities of these two events.

**Fig 2.**
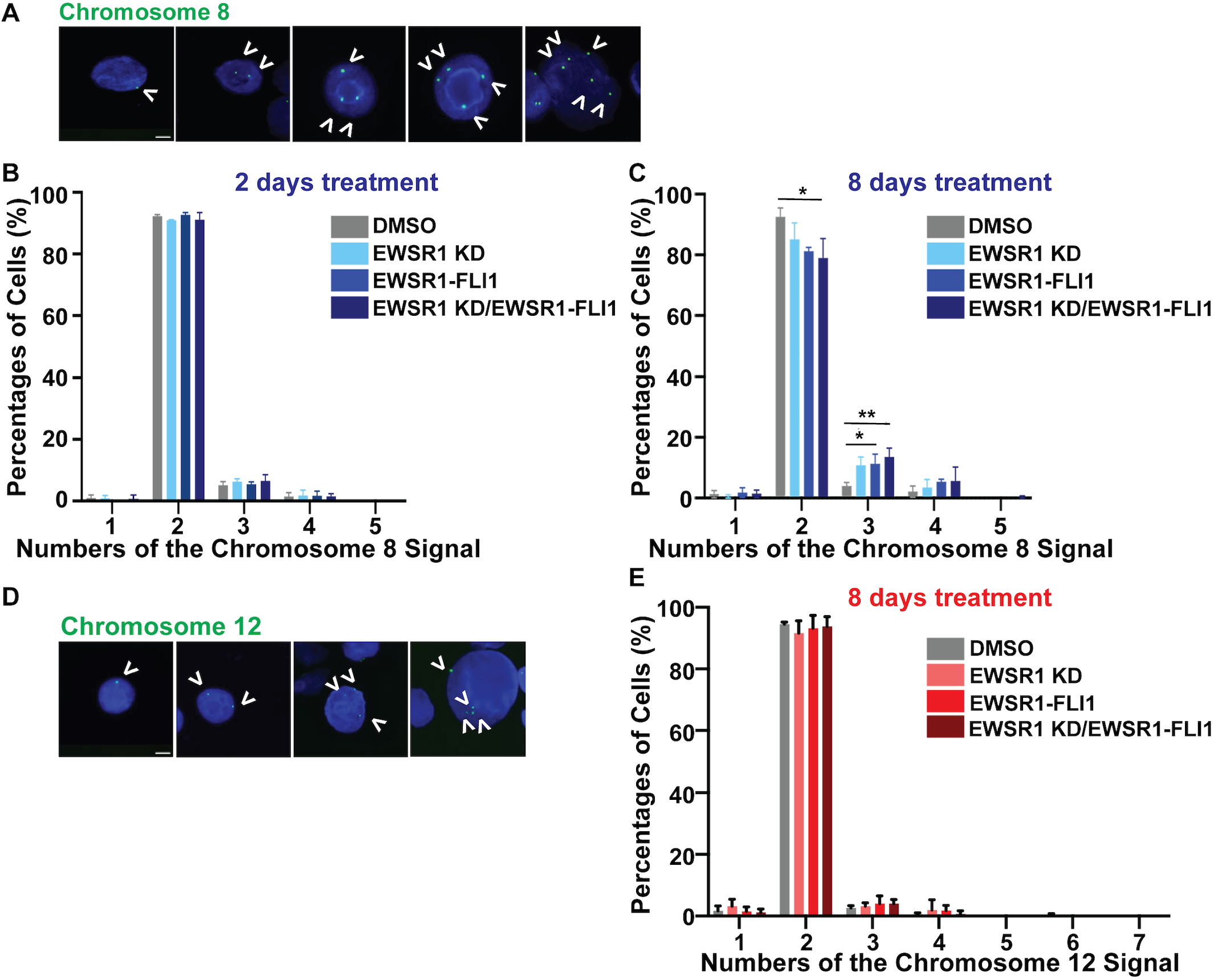
The combination of EWSR1 knockdown and EWSR1-FLI1 induces trisomy 8 but not trisomy 12 within eight days. **A**. Representative images show different numbers of chromosome 8 (Green, indicated with white arrow head), obtained from the AUX/DOX cells, visualized by FISH assays using the probe for chromosome 8. **B and C**. The (*AID-EWSR1/wt;EWSR1-FLI1/wt*) cells were treated with DMSO (control), AUX (EWSR1-KD), DOX (EWSR1-FLI1), and AUX/DOX (EWSR1-KD/EWSR1-FLI1) for 2 days and 8 days. The percentage of trisomy 8-carrying cells visualized with FISH assays is shown in Fig 2B (2days) and 2C (8 days), respectively. **D**. Representative images for the nuclei (AUX/DOX treated cells) visualized with the FISH assay using chromosome 12 probe (Green signal, indicated with white arrow head). **E**. The percentage of trisomy 12-carrying cells for 8 days of treatment with DMSO, AUX, DOX, and AUX/DOX for 8 days. Scale bar: 20 μm. **: P<0.01. Statistical analysis was done with one-way ANOVA, followed by Tukey’s multiple comparison test. Note that the samples without the * do not have any significant differences.

Since trisomy 12 is the second most frequently observed aneuploidy in Ewing sarcoma, we also tested the frequency of trisomy 12 induction using the same experimental conditions for FISH. To our surprise, the induction of one *EWSR1* loss and *EWSR1-FLI1* expression for eight days did not cause changes in the ploidy of chromosome 12, unlike the case of chromosome 8 (**Fig 2D and 2E**). Together, the one allele-derived EWSR1 and EWSR1-FLI1 expression cooperate to promote trisomy 8, but not trisomy 12, on day eight.

### The EWSR1-FLI1 interacts with EWSR1 during mitosis

Our previous study showed that EWSR1-FLI1 inhibits the activity of EWSR1 and induces mitotic dysfunction through physical interactions (*33*). Therefore, it is plausible that trisomy 8 is induced by the compromised activity of EWSR1 through two mechanisms. The first mechanism is the haploinsufficiency of *EWSR1* caused by losing one *EWSR1* allele due to the formation of *EWSR1-FLI1* fusion gene. The second mechanism is the dominant inhibition of EWSR1 by EWSR1-FLI1, which occurs through physical interaction. Since it was unknown whether the EWSR1-FLI1 interacts with EWSR1 during mitosis, we conducted Proximity Ligation Assays (PLA) in the (*AID-EWSR1/wt;EWSR1-FLI1/wt*) cells. The cells were treated with DMSO and DOX, and subjected to a PLA assay using an anti-FLAG antibody to recognize the AID-EWSR1 and an anti-mCherry antibody to recognize EWSR1-FLI1-mCherry. The EWSR1-FLI1 expressing cells (DOX treated cells) exhibited high intensity of PLA signals during prophase, metaphase and anaphase when compared to the control (DMSO treated) cells (**Fig 3A**). The intensity of PLA signals from three experiments reveled that the PLA signals of the DOX treated samples are significantly higher than the control (DMSO treated) cells (**Fig 3B**). The data suggests that EWSR1-FLI1 interacts with EWSR1 during prophase, metaphase and anaphase.

**Fig 3.**
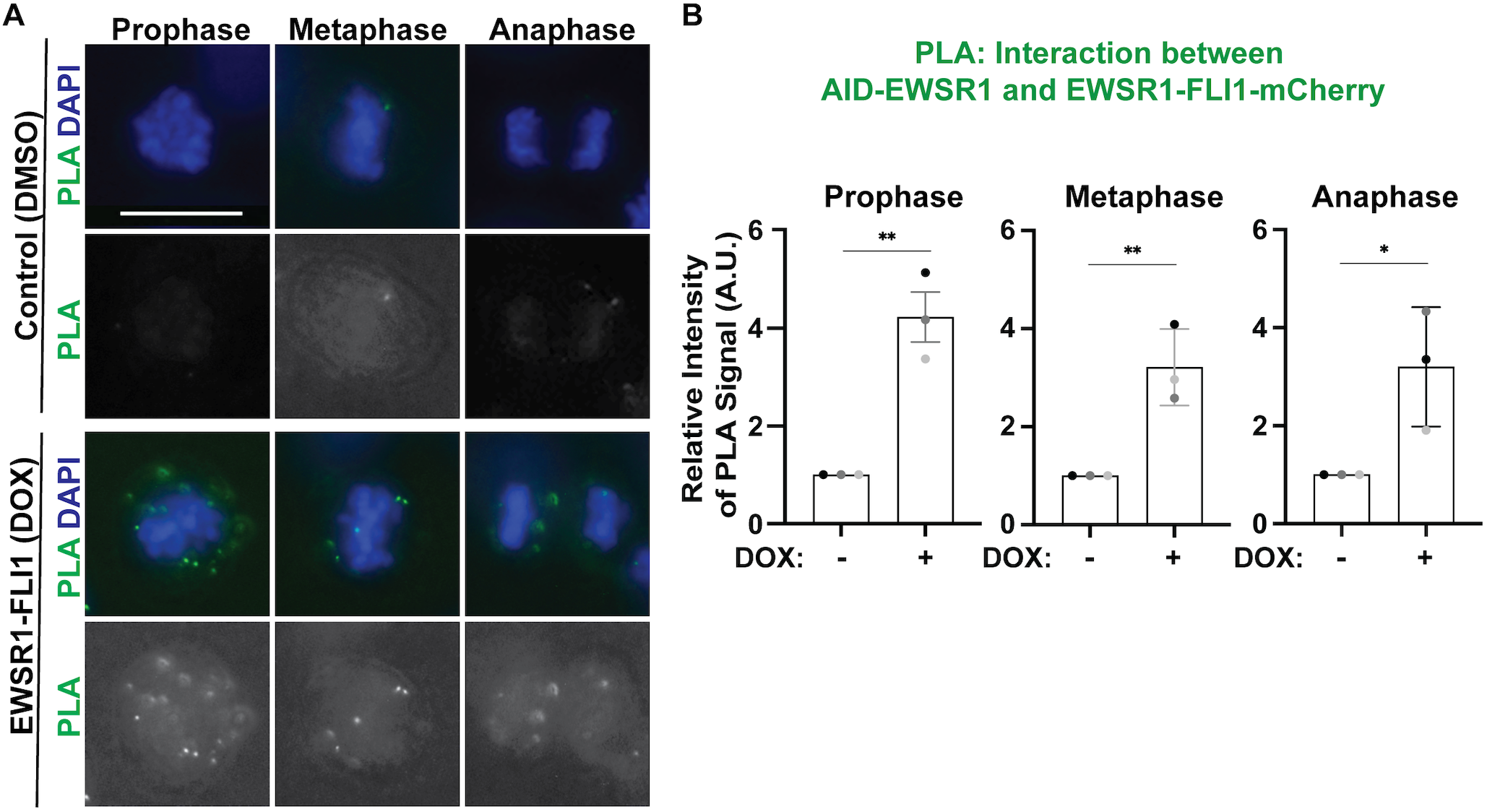
The EWSR1-FLI1 interacts with EWSR1 during mitosis. **A**. Representative images of (*AID-EWSR1/wt;EWSR1FLI1/wt*) cells at prophase, metaphase and anaphase were visualized by proximity ligation assay (PLA) after 4 days of treatment with control (DMSO) and EWSR1-FLI1-induced (DOX) cells. PLA was conducted using an anti-FLAG antibody for EWSR1 and an anti-mCherry antibody for EWSR1-FLI1. Scale bar: 20 μm. **B**. The intensity of PLA signals from the cells at mitosis (prophase, metaphase and anaphase) was quantified. **: P<0.01.

### Impaired interaction between EWSR1 and Aurora B leads to the induction of trisomy 8

The mesenchymal stem cell (MSC) has been suggested as the cell of origin of Ewing sarcoma (*41*). If the inhibition of EWSR1 activity is responsible for trisomy 8 induction, knockdown of EWSR1 in MSC should induce trisomy 8. We transfected EWSR1 siRNA and control siRNA into MSC for forty-eight hours, and conducted a FISH assay using a probe against chromosome 8. The immunocytochemistry confirmed the drastic knockdown of EWSR1 in the MSC transfected with EWSR1 siRNA compared to the control siRNA transfected MSC (**Fig 4A**). The FISH assay revealed that the incidence of trisomy 8 is significantly increased in the EWSR1 siRNA transfected MSC, whereas the control siRNA transfected cells did not show an increase in trisomy 8 (**Fig 4B**). The result suggests that EWSR1 knockdown promotes the induction of trisomy 8.

**Fig 4.**
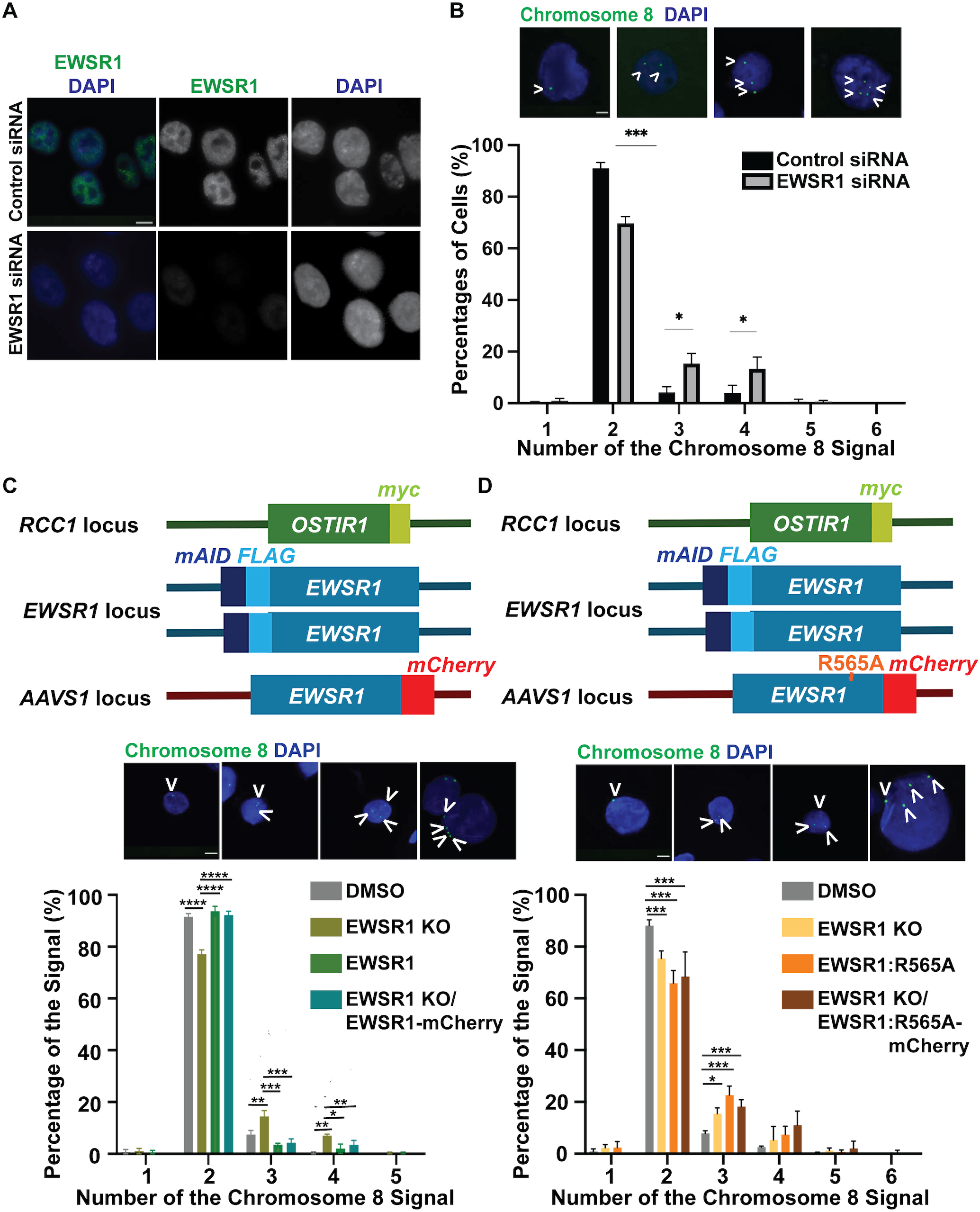
The re-expression of EWSR1 rescues the high incidence of trisomy 8 induced by EWSR1 knockout, whereas the EWSR1:R565A mutant cannot rescue the phenotype. **A**. MSC were transfected with Control siRNA and EWSR1 siRNA for forty-eight hours. **B**. Control and EWSR1 siRNA transfected MSCs were subjected to FISH assay using a probe for chromosome 8. **C**. The (*AID-EWSR1/AID-EWSR1:EWSR1-mCherry*) DLD-1 cells were treated with DMSO (control), AUX (EWSR1 knockout), DOX (EWSR1-mCherry overexpression) and AUX/DOX (EWSR1 knockout/EWSR1-mCherry overexpression) for 8 days, followed by FISH assay using a probe for chromosome 8. **D**. The (*AID-EWSR1/AID-EWSR1:EWSR1:R565A-mCherry*) DLD-1 cells were treated with DMSO (control), AUX (EWSR1 knockout), DOX (EWSR1:R565A-mCherry overexpression) and AUX/DOX (EWSR1 knockout/EWSR1:R565A-mCherry overexpression) for 8 days, followed by the FISH assay using a probe for chromosome 8. Scale bar: 20μm. *: P<0.05, and ***: P<0.001. One-way ANOVA, followed by Tukey’s multiple comparison test, was applied for the analysis.

To further highlight the importance of the loss of EWSR1 activity in inducing the trisomy 8, we utilized the (*AID-EWSR1/AID-EWSR1;EWSR1-mCherry/wt*) DLD-1 cell line. The cell line facilitates the degradation of EWSR1 from both alleles with AUX (EWSR1 knockout: KO) since both *EWSR1* alleles are tagged with mAID. The cell line also permits a rescue experiment by overexpressing EWSR1-mCherry with DOX within AUX-induced EWSR1 knockout background (**Fig 4C**). The cells were treated with DMSO, AUX, DOX, and AUX/DOX for eight days. Then the numbers of the chromosome 8 were visualized by FISH assay, and plotted their rates (**Fig 4C**). There was a significant increase in trisomy 8 and tetrasomy 8 in both allele-derived KO cells (AUX) compared to the control (DMSO) cells. However, no significant changes were observed in the aneuploidy of chromosome 8 in the AUX/DOX treated cells compared to the control cells. The results suggest that the EWSR1 KO derived from two *AID-EWSR1* alleles increases the incidence of trisomy 8 and tetrasomy 8, which is rescued by the concomitant expression of EWSR1-mCherry (**Fig 4C**).

To address whether the induction of trisomy 8 in EWSR1 KO cells is caused by the absence of interaction between EWSR1 and Aurora B kinase, we conducted an additional experiment using the (*AID-EWSR1/AID-EWSR1;EWSR1:R565A-mCherry/wt*) DLD-1 cell line. The cell line enables the knockdown of EWSR1 and the expression of the EWSR1:R565A mutant, which lacks the ability to interact with Aurora B (*22*). Treating the cells with AUX results in EWSR1 KO, while treating the same cells with DOX induces the expression of EWSR1:R565A-mCherry (**Fig 4D**). The expression of EWSR1:R565A (DOX) in the EWSR1 KO cells (AUX) failed to rescue the high incidence of trisomy 8, unlike wildtype EWSR1-mCherry. These results suggest that the interaction between EWSR1 and Aurora B is required for preventing the induction of trisomy 8 (**Fig 4D**). Here, we demonstrate that EWSR1-FLI1 collaborates with the one allele-derived EWSR1, leading to the induction of trisomy 8. This is likely caused by the dominant-negative inhibition of EWSR1 activity by EWSR1-FLI1 (**Graphic Abstract**). This is the first demonstration of the mechanism of the trisomy 8 induction.

## Discussion

The establishment of the (*AID-EWSR1/wt:EWSR1-FLI1/wt*) cell line has enabled us to systematically study aberrations that mimic the early stage of cells induced by the loss of *EWSR1* allele and *EWSR1-FLI1* expression. We discovered that high potential for colony formation and a high incidence of trisomy 8 are induced by the combination of one allele-derived EWSR1 and/or EWSR1-FLI1 expression. This condition appears to generate two inhibitory mechanisms of EWSR1. One is the haploinsufficiency of EWSR1, and the other is the dependent inhibition of EWSR1 activity by EWSR1-FLI1. One supportive observation is that the EWSR1-FLI1 and EWSR1 interact with each other. Second supportive observation is that EWSR1 knockdown by Auxin treatment in the (*AID-EWSR1/AID-EWSR1;EWSR1-mCHERRY/wt*) cell line was sufficient to induce a high incidence of trisomy 8. Conditional expression of EWSR1 rescued the high incidence of trisomy 8, whereas the EWSR1:R565A (Aurora B binding mutant) in the (*AID-EWSR1/AID-EWSR1;EWSR1:R565A-mCHERRY/wt*) cell did not. This result suggests that the interaction between EWSR1 and Aurora B prevents the induction of trisomy 8.

A novelty of this study is the collaborative activity of one allele-derived EWSR1 and EWSR1-FLI1 expression in inducing trisomy 8. To our surprise, we did not observe trisomy 12 within the same experimental condition. This data raises the important question of why there is a difference in induction duration between trisomy 8 and trisomy 12, and whether these aneuploidies contribute to the heterogeneity of Ewing sarcoma. One possible explanation for this question is that aneuploidies are induced at an equal frequency among all chromosomes, but the cells may have a proliferation advantage compared to cells carrying other aneuploidies. This is likely caused by a change in protein expression resulting from the acquisition of chromosome 8. For example, Rad21 is encoded on chromosome 8, and cells carrying trisomy 8 have been reported to mitigate replication stress due to the overexpression of the protein (*16*). The *c-MYC* gene, also encoded on chromosome 8, is a strong candidate for promoting the survival and proliferation of these cells. Another possibility is that the trisomy 8 is induced at a higher incidence than other trisomy carrying cells when it undergoes loss of one EWSR1 and EWSR1-FLI1 expression. If this is the case, the centromeres of chromosome 8 may have a specific condition that makes them more susceptible to trisomy 8 induction in the presence of the loss of one *EWSR1* allele and EWSR1-FLI1 expression.

To further illustrate the molecular mechanism of trisomy 8 induction, it is crucial to elucidate the role of the EWSR1 protein in aneuploidy induction for two reasons. Firstly, EWSR1 regulates faithful mitosis to ensure the accurate segregation of chromosomes, leading to an equal distribution of chromosomes in two daughter cells. One known cause of aneuploidy induction is merotelic attachment, where K-fibers nucleated from both centrioles attach to the kinetochore of one chromosome during metaphase. Our previous study demonstrated that EWSR1 interacts with and regulates Aurora B, a key kinase responsible for ensuring proper attachment of K-fibers to the kinetochore (*24*). The EWSR1:R565A mutant, lacking the ability to bind Aurora B kinase, is unable to reduce the high incidence of trisomy 8 in EWSR1 knockdown (*AID-EWSR1/AID-EWSR1*) cells, suggesting that EWSR1-dependent regulation of Aurora B plays a crucial role in preventing the onset of trisomy 8. It is essential to study the role of EWSR1 at the centromere and proximal-kinetochore (PKC) regions, especially to determine if its activity is compromised in chromosome 8 in Ewing sarcoma cells. Another important reason to elucidate the function of EWSR1 is its potential regulation of tubulin dynamics. EWSR1 is localized on mitotic spindles and is known to regulate acetylation of tubulins through interaction with HDAC6 (*42*). An important example of this is the correction of erroneous K-fiber-kinetochore attachment to ensure faithful mitosis. The dynamic stability of spindles is crucial for the process. To fully understand the role of EWSR1 in mitosis, it is important to understand how EWSR1 coordinates the activities of Aurora B and the dynamics of mitotic spindles. One possible model suggests that EWSR1 undergoes phase separation at the centromere-KPC region, providing a platform to coordinate the activity of Aurora B and modulate spindle dynamics at the proximal kinetochore sites simultaneously. Once we have acquired knowledge of this mechanism, it is important to investigate whether the EWSR1-FLI1 is a part of the complex with the EWSR1-Aurora B dimer/complex. To gain knowledge of this mechanism, it is important to elucidate the dynamics of the complex status of EWSR1 and Aurora B, and its localization throughout the mitotic stages.

A high incidence of trisomy 8 is observed in various types of cancers, including clear cell sarcoma, rhabdomyosarcoma, and myxoid liposarcoma (*43-45*). While these diseases do not express EWSR1-FLI1, they all carry a fusion gene that contains N-terminus of EWSR1 (e.g. EWSR1-ATF1 in clear cell sarcoma, EWSR1-DUX4 in rhabdomyosarcoma, and EWSR1-DDIT3 in myxoid liposarcoma (*46-48*). Therefore, it is important to investigate whether EWSR1-fusion expressing sarcomas induce trisomy 8 through the same mechanism as observed in this study. The importance of understanding the activities of trisomy 8 is underscored by the observation that patients with these cancer types typically have a poor prognosis. The gap in knowledge lies in the uncertainty about whether the trisomy 8-carrying cell itself is the drug-resistant cell, or if the daughter cells of the trisomy 8-carrying cell relapse or metastasize. In addition, it remains unclear whether trisomy 8-carrying cells affect the selection of other chromosomes. Therefore, the dynamics of trisomy 8 after cell division should be analyzed in future studies.

## Materials and Method

### Establishment and Maintenance of Cell Lines

In a previous study, we reported a EWSR1-knockdown (AID-EWSR1/AID-EWSR1) DLD-1 cell line. In this study, the (*AID-EWSR1/wt;EWSR1-FLI1/wt*) DLD-1 cell line was further generated by integrating the Tet-on EWSR1-FLI1 construct and guide RNA for the safe-harbor *AAVS-1* locus. The DLD-1 cell lines (*AID-EWSR1/WT*;*EWSR1-FLI1-mCHERRY/wt*), (*AID-EWSR1/AID-EWSR1*;*EWSR1-mCHERRY/wt), and* (*AID-EWSR1/AID-EWSR1;EWSR1:R565A-mCHERRY/wt*) were maintained in the McCoy medium with 10% FBS containing 1 μg/mL Blasticidin (Invivogen, #ant-bl), 200 μg/mL Hygromycin B Gold (Invivogen, #ant-hg), and Puromycin (Invivogen, #ant-pr) (2 μg/mL). The cells were treated with Auxin (500 μM) to induce the degradation of AID-EWSR1, and Doxycycline (1 μg/mL) to induce the expression of *EWSR1-FLI1-mCHERRY/wt, EWSR1-mCHERRY/wt*, and EWSR1:*R565A-mCHERRY/wt* for 2 to 8 days (*49*).

### Cell Growth Assay

The (*AID-EWSR1/wt;EWSR1-FLI1-mCHERRY/wt*) DLD-1 cells were seeded at 20% confluency (63,000 cells) in a 12-well plate and treated with DMSO, Auxin (500 μM), and/or Doxycycline (1 μg/mL). The number of cells was counted after forty-eight hours of treatment. One-fourth of the cells were added back to the plate and treated with the drug to count the cells again after another forty-eight hours.

### Colony Formation Assay

The cells were synchronized to mitosis using the conventional thymidine/nocodazole protocol, while treated with DMSO, AUX and/or DOX simultaneously. The cells were plated on a 6-well plate, and allowed to grow to form colonies for 14 days. Afterwards, the colonies were washed three times with 1X PBS and stained using crystal violet (C0775-100G, Millipore Sigma). The colonies were photographed with the iBright FL1000 imaging system (Thermo Fisher, Waltham, MA, USA).

### Western Blotting

Whole cell lysates obtained from cells treated with DMSO, AUX and/or DOX were subjected to western blotting. Primary antibodies used in this study were: mouse anti-C9 EWS antibody (1:1,000 dilution) (Santa Cruz, #sc-48404). Secondary antibodies used in this study are: IRDye 680RD donkey anti-mouse IgG (1:10,000 dilution, LI-COR, #926–68072). All images of western blotting were captured and quantified by LI-COR Odyssey Imaging System.

### Immunocytochemistry

The cells were seeded at 20% confluency on cover slips (Electron Microscopy Sciences. #72290-04), treated with DMSO, AUX and/or DOX for twenty-four hrs. The cells were fixed using 4% paraformaldehyde for 10 minutes at room temperature, washed with phosphate buffered saline (PBS) three times for 5 minutes each. The cells were permeabilized with ice-cold methanol for 10 minutes, followed by three washes with PBS for 5 minutes each at room temperature, followed by a blocking procedure using 1% fetal bovine serum in (FBS)/PBS for one hour. Then, the cells were treated with primary antibodies as follows: mouse anti-FLAG antibody (1:500 dilution) (Sigma, F3165); rabbit anti-mCherry antibody (1:500 dilution) (ROCKLAND, 600-401-P16); mouse anti-α-tubulin antibody (1:2,000 dilution) (Sigma, #T6074). After the wash with PBS three times, cells were subjected to the treatment with secondary antibodies: anti-mouse Alexa fluor 488 (1:1000 dilution) (Invitrogen,# A-11004) or anti-rabbit Alexa fluor 594 (1:500 dilution) (Invitrogen, #A32790). The cells were washed with PBS three times, and mounted on a slide with mounting media containing DAPI (Vectashield, H-1200).

### Fluorescence In Situ Hybridization (FISH) Assay

The cells were seeded in a 3.5 cm culture dish and treated with DMSO, AUX and/or DOX for 2 or 8 days. After trypsinization, the cells were centrifuged at 500 g for 5 minutes to collect the cell pellet. The cells were incubated in a hypotonic solution (water: media; 1:1), 1 mL for 10 minutes at room temperature, and centrifuged at 500 g for 10 minutes. Freshly prepared 1 mL of cold methanol: glacial Acetic acid (3:1) solution was added dropwise to the cell pellet. The cell suspension was incubated on ice for 20 minutes and centrifuged at 10,000 g for 10 minutes. Then, the pellet was resuspended in 500 μL of ice-cold methanol: glacial acetic acid (3:1) and incubated on ice for 10 minutes. The cell pellet was obtained by centrifuging at 10,000 g for 10 minutes. The pellet was finally dissolved in 200 μL ice-cold methanol: glacial acetic acid (3:1). The cell suspension was dropped onto a glass slide.

The slide with chromosome spread was incubated in a pre-warmed pretreatment solution (1% NP-40, 2× SSC, pH 7.0) at 37 °C for 30 min. The air-dried slide was dehydrated in 70%, 80%, and 100% ethanol for 2 minutes each. The prepared FISH probe solution (mixture of probe 2 μL and buffer 8 μL) was added onto the fixed cells on the slide, covered with a coverslip and sealed with masking tape, and placed on a 73 °C hot plate for 3 minutes in the dark. The slide was incubated at 37 °C in a humid chamber overnight. The tape and coverslip were removed and incubated the slide in a slide jar with the pre-warmed wash solution 1 (0.5xSSC, 0.1% NP40) at 73°C for 1.5 minutes and gently agitated the slides during the incubation. The slide was transferred to wash solution 2 (2x SSC, 0.1% NP40) at room temperature for 1.5 minutes. Finally, rinsed with distilled water and air-dried. The cells were mounted with the mounting media containing DAPI (Vectashield, H-1200) (*50*).

### Proximity Ligation Assay (PLA)

The (*AID-EWSR1/wt*;*EWSR1-FLI1-mCHERRY/wt*) DLD-1 cell lines was seeded at 20% confluency on cover slips (Electron Microscopy Sciences. #72290-04), treated with DMSO, AUX and/or DOX for four days. The cells were synchronized to the G2/M phase border by treating them with 5μM RO-3306 (CDK1 inhibitor, SML0569-5MG, SIGMA) for 20 hours, and released for 1 hour. The cells were fixed using 4% paraformaldehyde for 10 minutes at room temperature, washed with PBS three times for 5 minutes each. The cells were permeabilized by ice-cold methanol for 10 minutes, washed three times with PBS for 5 minutes each at room temperature. the remaining experimental procedures were conducted by following the protocol for the Duolink®proximity ligation assay (PLA®) kit (Sigma-Aldrich). The antibodies used in the experiments are listed below: mouse anti-FLAG antibody (1:250 dilution) (Sigma, F3165-1MG), rabbit anti-mCherry antibody (1:250 dilution) (ROCKLAND, 600-401-P16 (*51*).

### Gene knockdown using siRNA

Mesenchymal Stem Cells (MSC) were plated in 3 cm plates at 20% confluency. The cells were transfected with 480 pmol of EWSR1 siRNA (sc-35347) or control siRNA (sc-37007, Santa Cruz Biotechnology, Inc.) in 1 ml of medium following the manufacturer’s protocol (Santa Cruz Biotechnology, Inc.). Cells were cultured for forty eight hours, and then subjected to immunocytochemistry.

### Statistical Analysis

Graphs in this study are presented with the mean and standard deviation (S.D.) or standard error of the mean (S.E.M). Statistical analysis was conducted with one-way ANOVA followed by Tukey’s multiple comparison test, or two-tailed unpaired t-test using GraphPad9 software. The confidence of assays was defined at *p* < 0.05.

## Conflict of Interest

The authors declare that the research was conducted in the absence of any commercial or financial relationships that could be construed as a potential conflict of interest.

## Funding

The study was supported by the Braden’s Hope Foundation Pilot Grant, Massman’s family research fund, Alan B Slifka Foundation Research Fund, University of Kansas (KU) General Research Fund (GRF), and the National Cancer Institute (NCI) of the NIH under award number P30 CA168524. The content is solely the responsibility of the authors and does not necessarily represent the official views of the National Institutes of Health.

## Acknowledgement

We would like to thank Dr. Kristi Neufeld for providing support with image documentation.

---

**Fig. S1.**
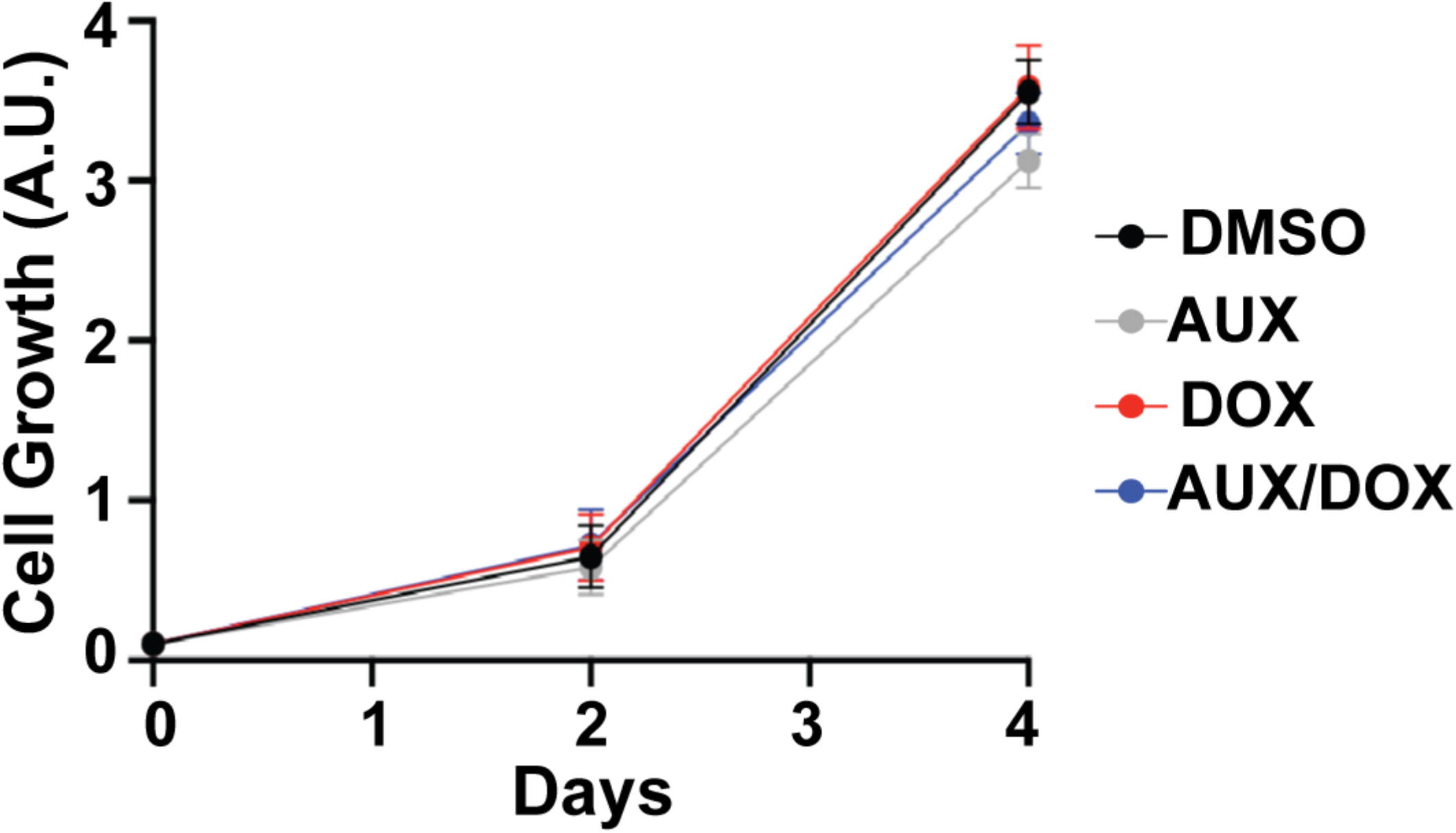

## Notes

### Competing Interest Statement

The authors have declared no competing interest.

